# Goal-directed action transiently depends on action space

**DOI:** 10.1101/783308

**Authors:** Laura A. Bradfield, Beatrice K. Leung, Susan Boldt, Bernard W. Balleine

**Affiliations:** Centre for Neuroscience and Regenerative Medicine, Faculty of Science, Faculty of Science, University of Technology Sydney, Sydney, Australia; St Vincent’s Centre for Applied Medical Research, Sydney, Australia; School of Psychology, University of New South Wales, Australia; School of Psychology, University of Sydney, Australia

## Abstract

Rats use spatiotemporal features of the environment to navigate to a goal, but whether representations of ‘action space’ are necessary for non-navigational goal-directed actions is unknown. We addressed this question by assessing goal-directed action control across contexts and under hippocampal inactivation and found that such actions do indeed rely on a representation of action space but only immediately after initial acquisition.

**One Sentence Summary:** Goal-directed actions depend on a hippocampal representation of action space immediately after initial encoding but not after a delay.

---

Goal-directed actions do not occur in a vacuum and yet how such actions rely on ‘action space’ – i.e. the spatiotemporal features of the environment in which they are acquired – is unknown. Some evidence suggests that goal-directed navigation relies on cognitive representations or ‘maps’ of the environment ^1,2^, but whether non-navigational actions also rely on such representations is far from clear.

The few rodent studies that have investigated this question appear to rule out a role for representations of action space in non-navigational goal-directed actions. For instance, goal-directed actions have been found to transfer fully across contexts^3^, and to be intact despite lesions of the dorsal hippocampus^4,5^; broadly considered the neural correlate of spatial cognitive maps in rodents ^6^. In contrast, several human studies demonstrate a central role for hippocampus in the regulation of non-navigational goal-directed decisions ^7–9^. One potentially crucial difference between these studies is that rodents are often trained to perform goal-directed actions over multiple days, whereas human subjects are typically trained and tested in a single session on the same day ^8,9^. Therefore, in the experiments described here, we investigated whether non-navigational goal-directed actions do indeed rely on a cognitive representation of action space, but only transiently, immediately following initial learning.

We used outcome devaluation tests^10^ to determine whether actions were goal-directed (see supplemental methods). Rats were trained to press two levers for two unique outcomes: pellets and sucrose. After training, the value of one of these outcomes was reduced relative to the other using sensory specific satiety^10^ after which the rats were given a choice between the levers in extinction (i.e. in the absence of feedback from pellets and sucrose delivery). Typically, rats respond on the lever associated with the still-valued (i.e. non-prefed) outcome and avoid the devalued lever on test, demonstrating control by both the current value of the outcome and the action-outcome association in accord with definitions of goal-directed action^10,11^.

Our first series of experiments (Experiments 1-2) investigated whether goal-directed actions depend on action space by altering the physical context after different amounts of training or different temporal delays. Our second series (Experiments 3-4) assessed the role of the hippocampus in any dependency of performance on action space using chemogenetic inactivation.

For Experiments 1A and 1B animals received 6 days of lever press training. For Experiment 1A, however, both levers earned polycose (O3) over days 1-5, and only on day 6 did the left (A1) and right (A2) levers uniquely earn the pellet and sucrose outcomes (i.e., A1→O1 and A2→O2; counterbalanced, design in Figure 1A). For Experiment 1B, however, both levers earned pellets and sucrose across all 6 days of training (design in Figure 1B). After day 6, animals in both experiments were given an outcome devaluation test in either the training context (Group SAME) or in a different context (Group DIFFERENT), with the expectation that only animals in Experiment 1B would show intact goal-directed actions across both contexts. This was the observed result. There were no group differences during lever press acquisition across any stage of either experiment (all Fs < 1, Figure S1A, Figure 1B, and Figure 1E). On test, however, animals in Experiment 1A showed intact devaluation (Valued > Devalued) if tested in the same context, but impaired devaluation (Valued = Devalued) if tested in the different context (Figure 1C), as there was a significant group x devaluation interaction, F(1,23) = 4.55, p = .044, and simple effect for Group Same, F (1,23) = 7.076, p = .014, but not Group Different, F < 1. By contrast, animals in Experiment 1B showed intact devaluation regardless of test context; there was a significant main effect of devaluation, F(1,19) = 11.78, p = .003, but no interaction, F < 1 (Figure 1F).

**Fig 1.**
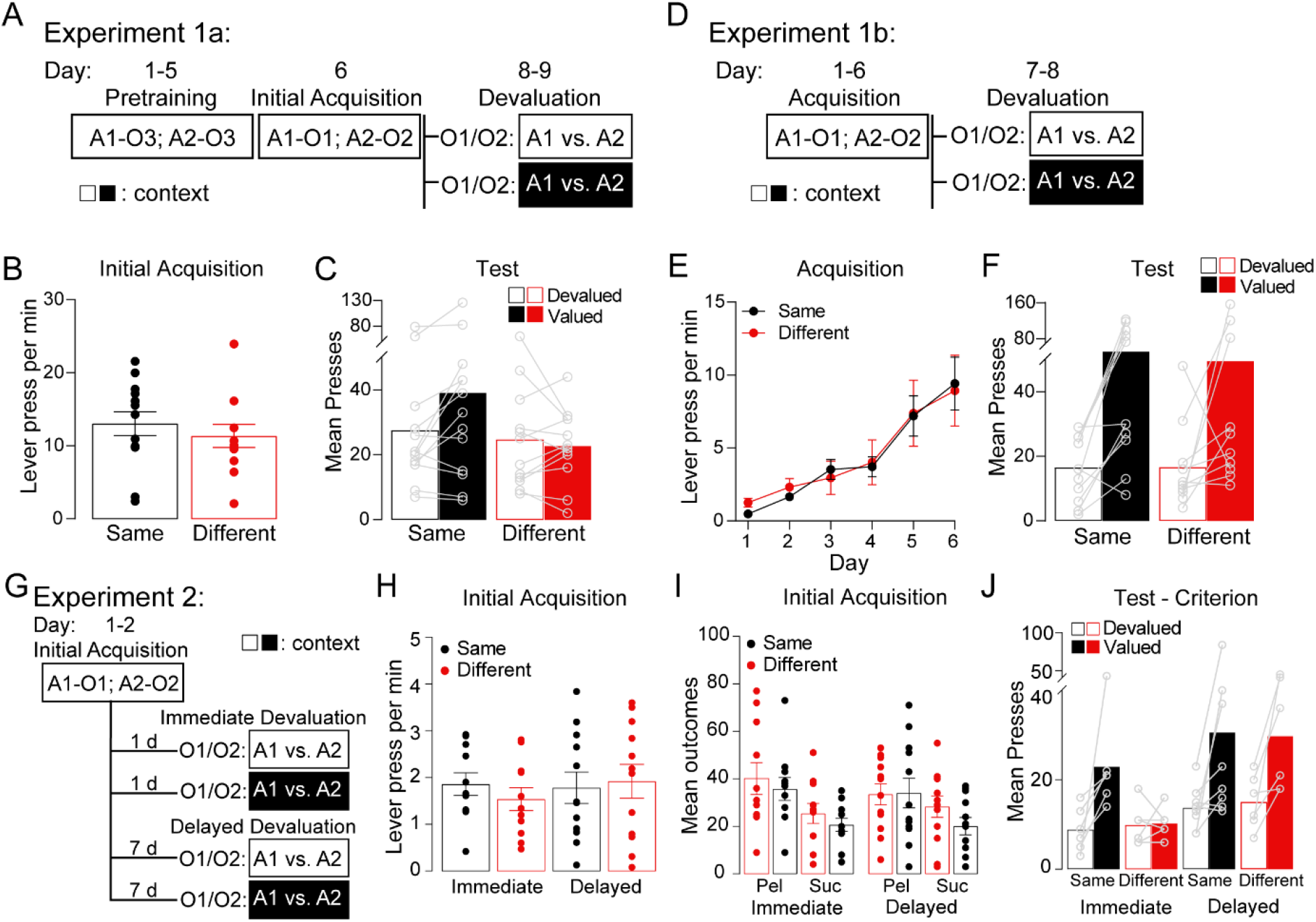
Outcome devaluation performance is impaired by a context switch immediately after limited training, but not when tested after additional training or after a delay. A) Design of Experiment 1A. Rats first were trained (Days 1-5) to press both left and right levers (Actions A1 and A2) for polycose (O3), then received a single day of training with each lever uniquely earning pellets or sucrose (O1 or O2, counterbalanced). Rats were given a specific satiety treatment for 1 hr on one or other outcome (O1 or O2) prior to a choice extinction test, A1 vs. A2. Group SAME were tested in the training context whereas group DIFFERENT were tested in a familiar, but different context. B) Lever presses per min on Day 6, C) Mean lever presses during devaluation test, D) Design of Experiment 1B. Rats were trained on the specific action-outcome contingencies (A1-O1, A2-O2) for all 6 days of training, then tested identically to rats in Experiment 1A). E) Lever presses per min during acquisition, F) Mean lever presses on test. G) Design of Experiment 2. All rats were trained on the specific contingencies (A1-O1, A2-O2) to a criterion of >20 action-outcome associations. Half of the animals received devaluation testing 1 day later in the SAME or DIFFERENT contexts (groups SAME-IMM, DIFF-IMM), and the other half were tested after a one week delay (groups SAME-DELAY, DIFF-DELAY), H) Lever presses per min during acquisition, I) Mean number of pellet (Pel) and sucrose (Suc) outcomes received during acquisition, J) Mean presses during devaluation test. Error bars represent ±1 SEM. See methods for details.

These experiments establish that goal-directed actions are initially context-dependent (Experiment 1A) but become context-independent with additional training (Experiment 1B). Experiment 2 (design in Figure 1G) sought to determine whether goal-directed actions become similarly independent of their physical context with the passage of time. All animals received 1-2 days of lever press training for the pellet and sucrose outcomes after which animals were tested in the same or different context either immediately (24 hours later, groups SAME-IMM and DIFF-IMM), or after a 1 week delay (groups SAME-DELAY and DIFF-DELAY). Again, lever pressing did not differ during acquisition (all Fs < 1, Figure 1H), nor did delivery of pellets/sucrose (Fs < 1, Figure 1I). On test, goal-directed action was context-dependent only when the devaluation test was immediate, and it was context-independent after a one-week delay (Figure 1J). Rats trained to criterion (> 20 outcomes on each lever, data for all rats is shown in Figure S1B) showed a significant group x lever interaction F(1,22) = 4.584, p = .044, supported by significant simple effects for Groups SAME-IMM, F(1,22) = 5.489, p = .029, SAME-DELAY, F(1,22) = 10.385, p = .004, and DIFF-DELAY F(1,22) = 6.018, p = .023, but not in Group DIFF-IMM, F < 1 (Figure 1J). These findings confirm that newly acquired goal-directed actions are transiently dependent on action space.

Next we sought to establish whether this transient dependency was reliant on hippocampal function, targeting dorsal hippocampus due to its well-documented role in spatial learning and memory in rodents^13^. In Experiment 3 (Design in Figure 2A) we used inhibitory hM4Di DREADDs to inactivate the CA1 region of dorsal hippocampus in one group and the CA2 region in a control group prior to devaluation testing, based on procedures validated previously^14,15^. Lever press acquisition was equivalent for all groups transfected with hM4Di DREADDs (all Fs < 1, Figure 2B), however group mCherry+CNO pressed significantly less than hM4Di+Veh, F(1,31) = 4.22, p = .049. Nevertheless, (for animals trained to criterion) devaluation was intact for group mCherry+CNO (Valued > Devalued), along with groups hM4Di+Veh, and hM4Di CA2+CNO. Only group hM4Di CA1+CNO was impaired (Figure 3C). There was a group (hM4Di CA1+CNO vs. the others) x devaluation interaction, F(1,31) = 5.19, p = .03, supported by significant simple effects for groups hM4Di+Veh, F(1,31) = 7.03, p = .012, mCherry+CNO, F(1,31) = 17.98, p = .00, and CA2+CNO, F(1,31) = 7.32, p = .011, but not group hM4Di CA1+CNO, F < 1.

**Fig 2.**
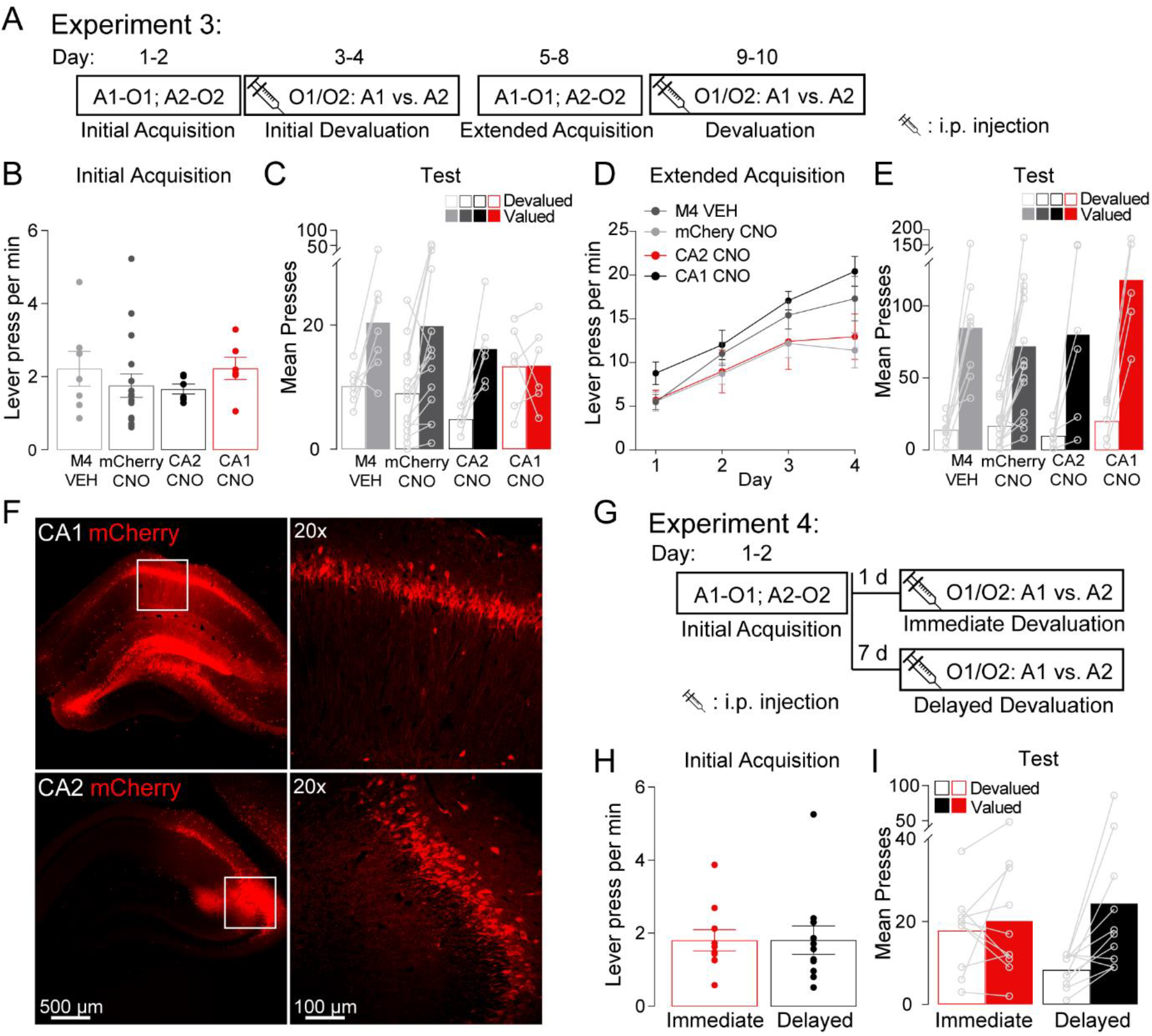
Chemogenetic inactivation of dorsal hippocampus (using hM4Di designer receptors exclusively activated by designer drugs: DREADDs) impaired devaluation performance when animals were tested immediately (1 day) after training, but not after additional training or a one week delay. A) Design of Experiment 3. All rats were trained on the specific pellet/sucrose contingencies (A1-O1, A2-O2) over 1-2 days to a criterion of ≥20 outcomes per lever. For test, animals were sated for 1 hr on one outcome (O1 or O2) prior to a choice test: A1 versus A2. 1 hr prior to test, animals received i.p. injections of either vehicle or clozapine-N-oxide (CNO) in accordance with their group assignment. Rats were then trained on the specific action-outcome contingencies for a further 4 days and tested again. CNO/Vehicle group assignment was altered for hM4Di-transfected animals between the first and second rounds of devaluation testing. B) Lever presses per min during initial acquisition (Days 1-2), C) Mean lever presses during the initial devaluation test (days 3-4), D) Lever presses per min during extended acquisition, E), Mean presses during the extended devaluation test (days 9-10), F) dorsal hippocampal neurons transfected with the AAV-hSyn-hM4D(Gi)-mCherry DREADDs virus (red) and close up of transfection in CA1 (top row), lateral CA2 region neurons transfected with the same virus, and close up of transfection in CA2 (bottom row), G) Design of Experiment 5. Rats were trained on contingencies A1-O1, A2-O2 over 1-2 days, then received devaluation tests either immediately (1 day later) or after a delay (1 week later). Rats received i.p. CNO injections 1 hr prior to test, H) Lever presses per min during acquisition, I) Mean lever presses on test. Error bars represent ±1 SEM.

Following this test, all groups were trained for a further 4-5 days and tested again. We re-arranged the group assignments after initial testing so that half of the animals that received CNO injections on test 1 received Vehicle injections on test 2, and vice versa. Once again, the hM4Di groups did not differ in lever pressing across days (largest F(1,31) = 1.76, p = .194), but group mCherry+CNO group pressed significantly less than group hM4Di+Veh, F(1,31) 6.16, p = .019, (Figure 3D). This time, however, all groups showed intact devaluation on test (Figure 3E). There was a main effect of devaluation, F(1,31) = 77.57, p = .00, which didn’t interact with any group differences, largest F(1,31) = 2.2, p = .148. This result shows that goal-directed actions are indeed initially hippocampally-dependent, but become hippocampally-independent over multiple days of training. This result is supported by supplemental experiments that employed muscimol infusions to inactivate dorsal hippocampus, Figure S2.

The final experiment (Experiment 4, design in Figure 2G) investigated whether goal-directed actions also become hippocampally-independent merely with the passage of time. For this experiment, the dorsal hippocampus (including CA1) was inactivated for all rats on test, half of which were tested immediately after training (1 day, i.e. hM4Di+CNO-IMM), and the other half tested after a one week delay (i.e. hM4Di+CNO-DELAY). Groups did not differ in their lever pressing during acquisition (F < 1, Figure 3H), but only rats tested after a delay showed intact devaluation on test (Valued > Devalued, F(1,19) = 8.86, p = .014), whereas those tested immediately did not, F < 1 (Figure 3I).

Collectively, these results support our central claim that goal-directed actions transiently depend on a cognitive representation of action space. If this representation is disrupted, either by alterations in context or by inactivation of dorsal hippocampus, the encoding of goal-directed actions is impaired. Our findings support this specific claim rather than a broader claim about contextual regulation of goal-directed actions, because Experiment 2 essentially involves two contextual alterations: physical (i.e. the change in physical context) and temporal (i.e. testing after immediately versus after a one week delay). Thus, if goal-directed actions were broadly context-dependent after initial training we would expect the delay alone to impair devaluation, even in the absence of an altered physical environment. In contrast to this prediction, however, our results found devaluation to be intact in both the same *and* different contexts after a delay (Figures 1J).

The effects that we have demonstrated here fit well with several of the known functions of hippocampus, such as systems consolidation ^17,18^ and episodic memory ^19,20^, and, for the first time, link these functions directly with decision-making involving choice between distinct courses of action based on changes in action value. The systems consolidation account^17,18^ suggests that the dorsal hippocampus (and specifically CA1) regulates short term memory formation and recall, which becomes hippocampally independent, possibly migrating to frontal cortical structures over the course of a week. The vast majority of the evidence for this theory has come from studies of conditioned reflexes in Pavlovian conditioning, particularly Pavlovian fear conditioning ^21,22^. In light of current findings, however, a link between systems consolidation and how this affects functionality in terms of action control warrants further consideration. Perhaps relatedly, our findings are also consistent with the theory that goal-directed actions initially rely on (contextually and hippocampally-dependent) episodic memories, but come to depend on context-free, extra-hippocampal, semantic memories over time^23,24^.

## Acknowledgments

We thank Genevra Hart for helpful discussions regarding these data. We thank Fred Westbrook for his feedback on the manuscript.

## Author contributions

L.A.B, B.K.L., and B.W.B designed the experiments, L.A.B., B.K.L., and S.B. performed the experiments, L.A.B, B.K.L. and B.W.B wrote the manuscript.

## Data and materials availability

Research data for this article (Figures 1–2 and S1-S3) is available for download at the following link: https://osf.io/xfmdp/?view_only=a12e3bdcb0164874be97579cf9ffbd8e.

