## Supplemental Materials for "Goal-directed action transiently depends on action space"

### Supplementary Materials

### Materials and Methods

All procedures were approved by the University of New South Wales Ethics Committee and/or or the University of Sydney Ethics Committee.

#### Subjects and Exclusions

Long-Evans Rats were housed in transparent amber plastic boxes ($0.5m^3$; 3-4 rats per box) located in a temperature- and humidity-controlled vivarium and were maintained on a 12 h light/dark cycle (lights on between 7:00 A.M. and 7:00 P.M.). Experiments 1a-1b, and S1a-S1b were conducted with male rats. All other experiments (2-4) were conducted using approx. 50% male and 50% female rats.

***Experiment 1a***
A total of 25 male rats, weighing between 350-500g at the beginning of the experiment were used as subjects, 13 in group SAME, and 12 in group DIFFERENT.

***Experiment 1b***
A total of 23 male rats, weighing between 350-500g at the beginning of the experiment were used as subjects. Two animals were excluded for failing to lever press, thus final experimental numbers were 11 rats in group SAME, and 10 rats in group DIFFERENT.

***Experiment 2***
A total of 46 rats (22 males and 24 females), males weighing between 350-500g at the beginning of the experiment, and females weighing between 200-300g at the beginning of the experiment were used as subjects. Experimental numbers were: 11 rats in group SAME-IMM, 11 rats in group DIFF-IMM, 12 rats in group SAME-DELAY, and 12 rats in group DIFF-DELAY.

Twenty animals were excluded from the reported test for not reaching criterion (> 20 outcomes per lever, although these animals were tested, and test data for all animals are shown in Figure S1B). Therefore, final experimental numbers were 6 rats in group SAME-IMM, 6 rats in group DIFF-IMM, 8 rats in group SAME-DELAY, and 6 rats in group DIFF-DELAY.

***Experiment 3***
A total of 55 rats (28 males and 27 females) were used as subjects. Males weighed between 350-500g and females weighed between 200-300g at the beginning of the experiment.
Four animals were excluded for misplaced DREADDs expression. Sixteen animals were excluded from the final test for not reaching criterion (> 20 outcomes per lever).

Experimental numbers for the first round of devaluation testing (after 1-2 days of training) were: 7 rats in group hM4Di+Veh, 16 rats in mCherry+CNO, 6 rats in hM4Di CA1+CNO, and 6 rats in hM4Di CA2+CNO.

For hM4Di-transfected animals, half of those that received CNO during the first round of testing were switched to Vehicle for the second round of testing (after 6 days of training), and vice versa. Experimental numbers for the second round of devaluation testing (after 6 days of training) were 8 rats in group hM4Di+Veh, 16 rats in mCherry+CNO, 5 rats in hM4Di CA1+CNO, and 6 rats in hM4Di CA2+CNO.

The remaining half received consistent injections prior to both tests.

***Experiment 4***
A total of 24 rats (12 males and 12 females) were used as subjects. Males weighed between 350-500g and females weighed between 200-300g at the beginning of the experiment.
One animal was excluded for an infection at the injection site. Two animals were excluded from the final test for not reaching criterion (> 20 outcomes per lever).
Experimental numbers on test were: 10 rats in hM4Di-IMM, and 11 rats in hM4Di-DELAY (N = 21).

***Experiment S1a***
A total of 24 male rats, weighing between 350-500g at the beginning of the experiment were used as subjects. Five rats were excluded from the final analysis due to cannula misplacement (n = 4) or infection (n = 1). Thus the final experiment numbers were 9 rats in group SALINE, and 10 rats in group MUSCIMOL.

***Experiment S1b***
A total of 19 male rats, weighing between 350-500g at the beginning of the experiment were used as subjects. Three rats were excluded from the final analysis due to cannula misplacement. Thus the final experiment numbers were 9 rats in group SALINE, and 7 rats in group MUSCIMOL.

**Apparatus**

For all behavioural experiments, training was conducted in 16 MED Associates operant chambers enclosed in sound- and light-attenuating cabinets. Each chamber was fitted with a pellet dispenser capable of delivering a 45 mg grain food pellet ( F0165, BioServ Biotechnologies), to a recessed magazine inside the chamber, as well as two pumps fitted with syringes outside the chamber, capable of delivering 0.2 mL of either 20% sucrose solution (white sugar, Coles) diluted in $H_2O$ or 20% maltodextrin solution (Poly-Joule, Nutrica) diluted in $H_2O$, each delivered to separate compartments of the recessed magazine inside the chamber. The chambers also contained two retractable levers that could be inserted individually on the left and right sides of the magazine. Head entries into the magazine were detected via an infrared photobeam. Unless otherwise stated, the operant chambers were fully illuminated during all experimental stages, illumination was provided by a 3W, 24V house light located on the upper edge of the wall opposite to the magazine. All training sessions were pre-programmed on two computers located in a separate room through the MED Associates software (Med-PC), these computers also recorded the experimental data from each session.

**Contexts**

Experiments 1a, 1b, and 2 employed two distinct contexts. One of these contexts constituted the bare, unadorned chamber with a paper towel placed in the bedding that had 0.5ml of 10% peppermint essence added. For the other context, laminated sheets of black and white vertical stripes were positioned on the transparent walls of the chambers, smooth Plexiglas sheets were placed on the floor, and a paper towel with 0.5 ml of 10% vanilla essence was placed in the bedding. Therefore, these contexts differed along visual (transparent vs. striped walls), tactile (grid vs. smooth floor), and olfactory (peppermint vs. vanilla) dimensions.

For these experiments (1a, 1b, and 2), animals received one magazine training session in each context during which both pellet and sucrose outcomes were delivered, one on each day (order counterbalanced), and two more pre-exposures to the ‘different’ context after lever press training sessions. This served to familiarise the animals to the different context and reduce neophobia. Pre-exposure sessions lasted 40 min during which no levers were extended and no food was delivered.

**Surgery**

Rats were anaesthetized with 3% inhalant isoflurane gas mixed with oxygen, delivered at a rate of 0.5 L/min throughout surgery. Anaesthetized rats were placed in a stereotaxic frame (Kopf Instruments). An incision was made into the scalp to expose the skull surface and the incisor bar was adjusted to place bregma and lambda in the same horizontal plane.

For Experiments 3 and 4, following the scalp incision a small hole was drilled into the skull above the hippocampus, either above the CA1 region (Half of the animals in Experiment 4, and all animals in Experiment 5, anteroposterior, -3.8, mediolateral, ±2.5, dorsoventral, -3.5 for males, and anteroposterior, -3.6, mediolateral, ±2.5, dorsoventral, -3.5 for females) or above the CA2 region (Experiment 4 only, anteroposterior, -3.8, mediolateral, ±2.5, dorsoventral, -3 for males, anteroposterior, -3.6, mediolateral, ±2.5, dorsoventral, -3 for females). A 1.0 µL glass syringe (Hamilton Company) connected to an infusion pump (Pump 11 Elite Nanomite, Harvard Apparatus) was lowered into the brain, and rats received 0.75µl infusions of either AAV8-hSyn-hM4D(Gi)-mCherry (Experiments 4 and 5) or AAV8-hSyn-mCherry (Experiment 4 only), at a rate of 0.15µL per minute. All DREADD viruses and control fluorophores were obtained from UNC Vector Core based on plasmids gifted by Bryan Roth and Karl Deisseroth. Following infusions, the needle was left in place for a further 2 min for diffusion before being retracted. The wound was subsequently closed with staples (Stoelting) and cleaned, after which rats were injected with a prophylactic (0.4 mL) dose of 300 mg/kg procaine penicillin (i.p), and 0.1mL of the analgesic Rymadil (s.c.). Rats were given a minimum of 3 weeks of recovery time following surgery to allow sufficient viral expression.

For Experiments S1a and S1b, a small hole was then drilled into the skull above the hippocampus (all co-ordinates in millimetres relative to bregma: anteroposterior, -3.8, mediolateral, ±3.2, dorsoventral, -2.5 at an angle of 15 degrees) in each hemisphere. A 26 gauge guide cannula (Plastics One) was then implanted into the hole, the tip of which was aimed towards the CA1 region of the hippocampus. Guide cannulas were maintained in position with dental cement, and dummy cannulas were kept in each guide at all times except during microinfusions. The wound was subsequently stapled and cleaned, after which rats were injected with a prophylactic (0.4 mL) dose of 300 mg/kg procaine penicillin interperitoneally (i.p), and 0.1mL of the analgesic Rymadil subcutaneously (s.c.). Rats were given one week of recovery following surgery during which they were weighed and monitored each day.

**Drug Infusions**

In Experiments S1a and S1b, rats received bilateral intra-hippocampal infusions of 5-aminomethyl-3-hydroxyisoxazole (muscimol, M1523, Sigma). A total of 0.5 µL of muscimol (0.5mg/ml) was injected at .32 µL/min. Control rats received a saline infusion at the same rate. Microinfusions were conducted using a 33 gauge infusion cannulas (Plastics One) that extended 1mm beyond the guide cannula. Infusion cannulas were inserted into the guide cannulas and connected to 25µL glass syringes (Hamilton Company) fitted on an infusion pump (PHD ULTRA 4400, Harvard apparatus). The infusion cannulas were left in place for 1 min following infusions to allow diffusion of the drug. Rats were placed back in their home cage for 20 min prior to behavioural training/testing to permit the drug to take effect.

**Drugs for i.p. injection**

Experiments 3 and 4. Clozapine-*N*-Oxide (CNO; National Institute of Mental Health Drug Supply Program) was dissolved 0.8% HCl in water to a concentration of 7 mg/mL, and the pH was adjusted to 4.5. A solution of 0.1% HCl in water of the same pH was used as vehicle. Drug or vehicle was injected i.p. 1 hr prior to the onset of instrumental testing (i.e. immediately prior to being placed in devaluation boxes), at a volume of 1 mL/kg, hence the dosage was 7 mg/kg. We have previously demonstrated the viability of this procedure, and its efficacy in reducing firing in hM4Di DREADDs-transfected cells in dorsal hippocampus using electrophysiology[1].

**Food restriction**

Rats underwent 3 d of food restriction prior to the onset of magazine training and this continued throughout the duration of the experiment. During this time, they received 5 g (for females) or 8 g (for males) of home chow daily for the first two days, and 7-12 g (females) and 10-15 g (males) from the third day until the end of the experiment. Their weight was monitored daily to ensure it remained above 85% of their pre-surgery body weight at all times.

**Behavioral Procedures**

Please note that we employed random ratio schedules during training [2] as well as choice tests [3] in all experiments to promote goal-directed behavior and prevent the transition to habitual responding, even after multiple days of training.

**Magazine training**

Rats in Experiments 1-2 were given two sessions of magazine training (one in each context), and rats in Experiments 3-4 and S1a-S1b were given one session of magazine training. For these sessions, the house light was turned on at the start of the session and turned off when the session was terminated. No levers were extended. For rats that received polycose pretraining first (Experiments 1a, S1a, and S1b), 30 deliveries of 20% polycose solution were delivered into the magazine on a random time 60 s schedule (RT60). For rats that only had lever press training for pellets and sucrose solution (Experiments 1b, 2, 3 and 4), 20 deliveries of pellets and 20 deliveries of 20% sucrose solution were delivered on independent RT60 schedules.

### Polycose Pretraining

Rats that had polycose pretraining were trained to press two levers that earned the same outcome polycose, prior to receiving lever press training for pellet and sucrose solution. Each session lasted for a maximum of 50 min and consisted of four periods where a single lever was inserted into the chamber (i.e. two periods for each lever) separated by a 2.5 min time out period in which the lever was retracted and the house light was turned off. Each period ended after 20 outcomes had been earned or 10 min elapsed. The order of presentation of each lever was pseudorandom.

On day 1, lever presses on each levers were continuously reinforced with a polycose solution. On days 2 and 3, the schedule of reinforcement shifted to a random ratio (RR) 5 schedule such that the probability of polycose delivery was .2 for each action. On days 4 and 5, the schedule of reinforcement shifted to a RR10 schedule such that the probability of a delivery of the outcome was 0.1 for each action.

### Lever press training for pellets and sucrose solution

Each session lasted for a maximum of 50 min and consisted of four periods where a single lever was inserted into the chamber (i.e. 2 periods for each lever) separated by a 2.5 min time out period in which the lever was retracted and the house light was turned off. Each period ended after 20 outcomes had been earned or 10 min elapsed. The order of presentation of each lever was pseudorandom.

Rats that had previously received polycose pretraining on days 1-5 (Experiments 1a, S1a, and S1b), received a single contingency training session on day 6 where lever presses earned sucrose solution and pellets on a RR10 schedule. For half of the animals in each group, the left lever earned pellets and the right lever earned a sucrose solution, and the remaining animals were trained on the opposite action-outcome contingencies (counterbalanced).

For rats that received 6 days of lever press training (Experiment 1b) lever presses were continually reinforced with pellets or sucrose solution on day 1. After, the schedule of reinforcement shifted to a RR5 schedule on days 2-3 and a RR10 schedule on days 4-6. Action-outcome contingencies were counterbalanced for each group.

For rats that received 1-2 days of lever press training (Experiments 2, 3, and 4), lever presses were initially continually reinforced with pellets or sucrose solution in the first period each lever was extended (i.e. first 10 min on each lever). If rats earned more than 10 outcomes on both levers in the first 25 mins, they were moved to a RR5 schedule. Rats that did not earn more than 10 outcomes on both levers remained on a continually reinforced schedule during the entire 50 min session. Animals that earned more than 20 outcomes per lever by the end of day 1 (i.e. animals that reached criterion) were not trained on day 2. Action-outcome contingencies were counterbalanced for each group.

### Outcome Devaluation Tests

During devaluation tests, rats were first placed in the devaluation chambers (which were in a separate room from the operant chambers) and provided with *ad libitum* access to one of the previously earned outcomes (pellets or sucrose solution) for 1 hr. After prefeeding, animals

were placed in the operant chambers and given a choice test with both levers available for 5 min but no outcomes were delivered. The following day, rats were given another devaluation test for which they were prefed the opposite outcome. That is, if previously prefed on pellets they were now prefed on sucrose solution, and vice versa.
All rats received one 1 hr pre-exposure session to the devaluation chambers following the final lever press training session, in which they were fed a little bit of their daily chow. This served to habituate animals to the devaluation chambers to reduce neophobia.

Each individual experiment employed a variant on the same procedures, as described below.

***Experiment 1a***
Rats first received polycose pretraining in one context (context alterations are described above). After, rats received a single session of lever press training on pellets and sucrose solution in that same context. The following day, rats received outcome devaluation tests where they were first placed in the devaluation chambers for prefeeding. Immediately after prefeeding, animals were placed in the operant chambers with the same or a different context to that in which they had been trained and given a choice test between levers. The following day, rats were given another devaluation test for which they were prefed the opposite outcome. Context assignment for test did not change between days, such that animals were either tested in the ‘same’ context on both test days, or the ‘different’ context on both test days (although context *identity* was counterbalanced, such that the ‘same’ context was the vanilla/stripey/smooth context for half of the rats, and the unadorned/grid/peppermint context for the rest, and likewise for the ‘different’ context).

***Experiment 1b***
Rats received lever training sessions for pellets and sucrose solution for 6 days. After, rats received outcome devaluation testing in the same of different context as described in Experiment 1a. Context assignment for test did not change between days, such that animals were either tested in the ‘same’ context on both test days (i.e. one with each outcome), or the ‘different’ context on both test days.

***Experiment 2***
Rats received lever press training for pellets and sucrose solution for 1-2 days.
Outcome devaluation was conducted identically to that described for Experiments 1a and 1b, except that half of the animals received the devaluation tests one day after the final lever press session (Immediate; day 3-4), whereas the other half received the devaluation tests one week after the final lever press session (Delayed; day 9-10). Again, context assignment did not change between days.

***Experiment 3***
Rats received lever press training for pellets and sucrose solution for 1-2 days. After, rats received outcome devaluation testing as described above (Initial Acquisition; day 3-4). Rats then received additional lever press training sessions (day 5-8) for pellets and sucrose solution. Following this extended training, rats received a second set of devaluation tests (Extended Acquisition; day 9-10).

Prior to each devaluation day, animals were subject to i.p. vehicle or CNO injections immediately before being placed into the devaluation chambers. Once in these chambers, prefeeding procedures took place as described before. After prefeeding, rats were immediately placed into the operant chambers and given a 5 min choice test between levers.

***Experiment 4***
Rats received lever press training for pellets and sucrose solution for 1-2 days. After, half of the animals received devaluation tests one day after the final lever press session (Immediately; day 3-4), and the remainder were tested one week later (Delayed; day 9-10). Prior to each devaluation day, animals were subject to i.p. vehicle or CNO injections immediately before being placed into the devaluation chambers.

***Experiments S1a***
Rats first received polycose pretraining for 5 days. On day 6, rats received an intra-dorsal hippocampal infusions of saline or muscimol as described above. Rats were then placed back into their home cages for 20 min for the drug to take effect. After 20 min, rats were placed into the operant chambers for a single contingency training session for pellets and sucrose solution. After, rats received outcome devaluation testing across two days (day 7-8) as described above.

***Experiment S1b***
Rats first received polycose training for 5 days followed by 1 day of lever press training for pellets and sucrose solution. After, rats received outcome devaluation testing as described above (day 7-8). Following prefeeding on each devaluation day, rats were immediately administered intra-dorsal hippocampal infusions of saline or muscimol. Rats were then placed back into their home cages for 20 min for the drug to take effect before being tested in the operant chambers.

**Statistical Analysis**

Data was analysed using planned orthogonal complex contrasts and follow up simple effects. The per-contrast error rate was controlled at $\alpha = .05$ for each contrast in the procedure described by Hays[4].

**Histology**

Rats were rapidly anaesthetized with sodium pentobarbital (300 mg/kg i.p., Virbac) and transcardially perfused with 400mL of 4% paraformaldehyde in 0.1 M sodium phosphate buffer (PB; pH 7.4).

Brains collected for cannula placement only were post-fixed for 1 h in the same fixative and placed in 20% sucrose in phosphate buffered saline (PBS pH 7.2) overnight. 40 μm coronal sections were cut using a cryostat (CM1520, Leica Microsystems). Every third section was collected on a slide and stained with cresyl violet. Slides were examined for placement of the cannula tip and evidence of infusion. Animals were excluded when the placement of the guide cannulas were misplaced or when there was an infection.

Brains collected for further immunofluorescence analysis to determine the location of viral infections were post-fixed overnight at 4°C. Coronal sections (30 μm) were collected with a vibratome (VT1000, Leica Microsystems) and stored at -20°C in a solution containing 30% ethylene glycol, 30% glycerol in PB, until they were rinsed four times for 10 min in PBS, mounted on slides and cover-slipped in Vectashield mounting medium (VEH1400, Vector laboratories). Images were obtained using an Olympus FV1000 confocal microscope. For each rat, sections were selected along the rostral-caudal axis of dorsal hippocampus. The extent of the infection was determined using the boundaries defined by Paxinos and Watson [5]. Animals were excluded when the infection were not in the boundaries of the targeted region, or when infection was minimal or not observed.

**Figures S1-S4**

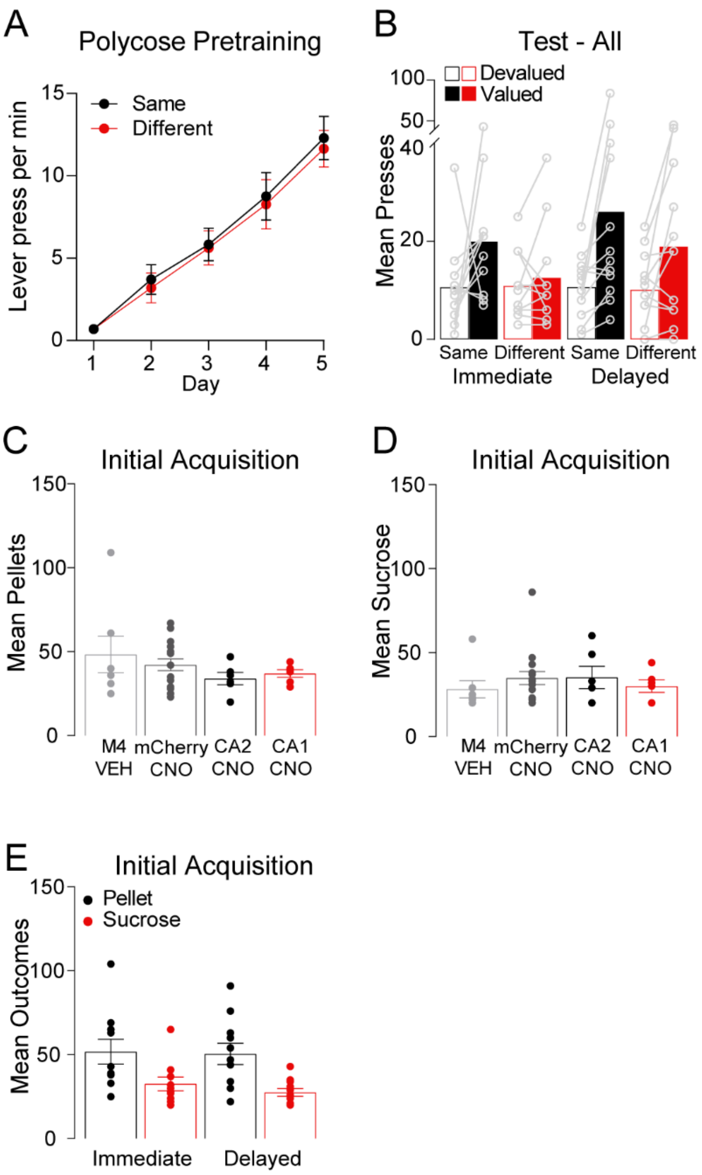


**Figure S1. Supplementary experimental data for Experiments 1-4.** A) Lever presses per min during polycose pretraining for Experiment 1a, B) Mean presses during devaluation test for Experiment 2 for all animals (i.e. not criterion animals only). For this test, Diff-Imm vs. other groups x devaluation interaction is $F(1,42) = 3.262$, $p = .078$. Simple effects (Valued > Devalued for all groups, $p < .05$, except for Diff-Imm, $F<1$). C) Mean pellet deliveries during initial training for Experiment 3, F) Mean sucrose deliveries for Experiment 3, G) Mean pellet and sucrose deliveries for Experiment 4.

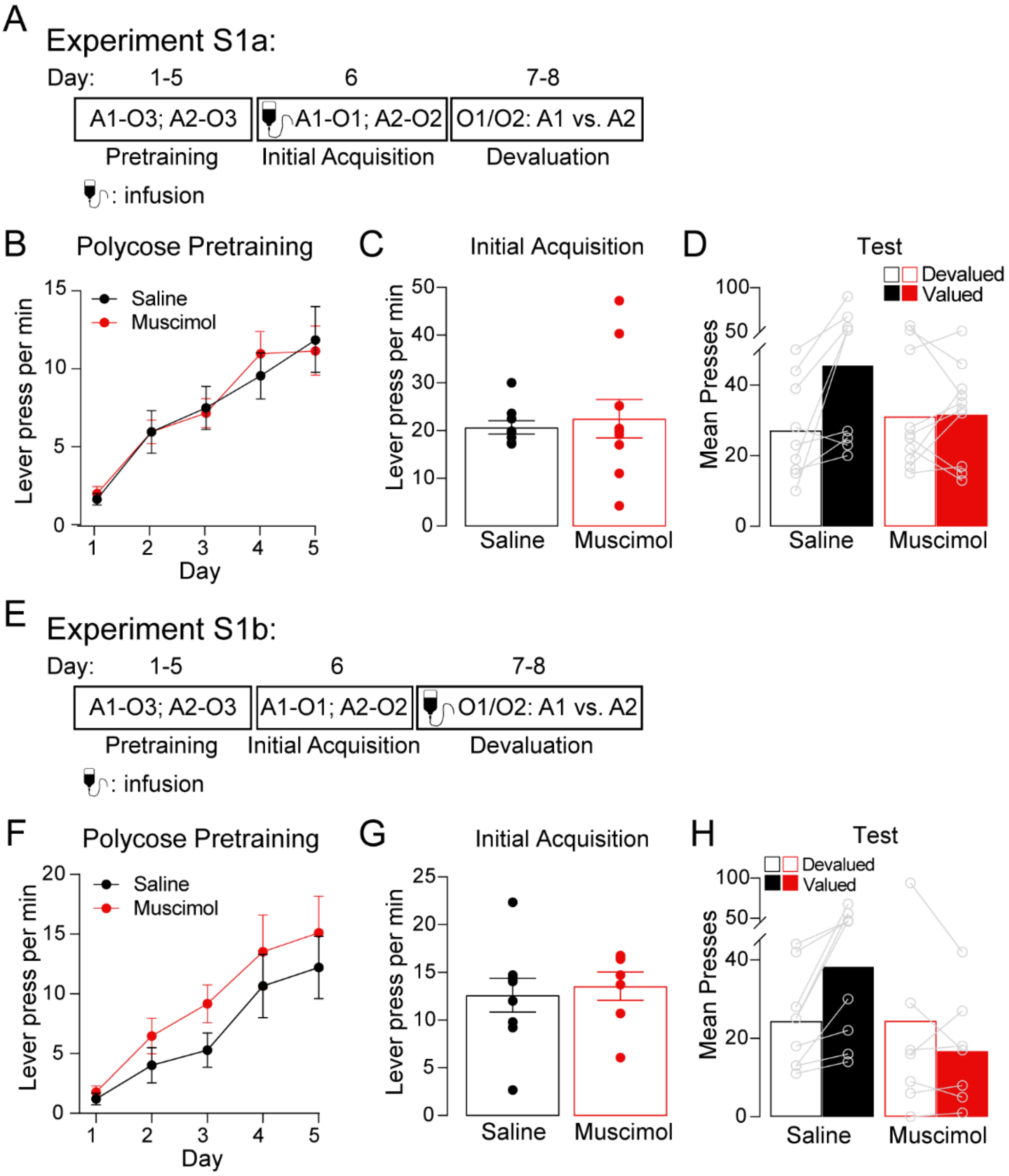


**Figure S2. Supplementary experiments S1a and S1b.** A) Design of Experiment S1a. Rats first were trained (Days 1-5) to press both left and right levers (A1 and A2) for polycose (O3), then received a single day of training with each lever uniquely earning pellets or sucrose (O1 or O2, counterbalanced). Prior to this single training session, half of the animals received control (saline) infusions into dorsal hippocampus, whereas the remaining animals received muscimol-induced inactivation. For test, rats were given a specific satiety treatment for 1 hr on one outcome (O1 or O2) prior to a choice test: A1 vs. A2, B) Lever presses per min during polycose pretraining, there were no group differences, $F<1$, a linear main effect $F(1,17) = 67.5$, $p = .00$, and no interaction, $F<1$, indicating that all rats acquired lever press responding, C) Lever presses per min on Day 6 of lever press training (pellets and sucrose delivered), responding did not differ between groups on this day, $F<1$, D) Mean lever presses during devaluation test. There was a group x devaluation interaction, $F(1,17) = 6.56$, $p = .02$, supported by a significant simple effect for group SAL, $F(1,17) = 13.47$, $p = .002$, but not group MUSC, $F<1$. E) Design of Experiment S1b. The design was identical to that for S1a, except that intra-hippocampal drug infusions took place 20 mins prior to devaluation testing (i.e. immediately after prefeeding), rather than prior to Day 6 training. F) Lever presses per min during polycose pretraining, there were no group differences, $F(1,14) = 1.02$, $p = .33$, G) Lever presses per min on Day 6 of lever press training (pellets and sucrose delivered), groups again did not differ on lever pressing, $F < 1$. H) Mean lever presses during devaluation test. There was a group x devaluation interaction, $F(1,14) = 6.09$, $p = .027$, supported by a significant simple effect for group SAL, $F(1,14) = 5.861$, $p = .03$, but not group MUSC, $F<1$.

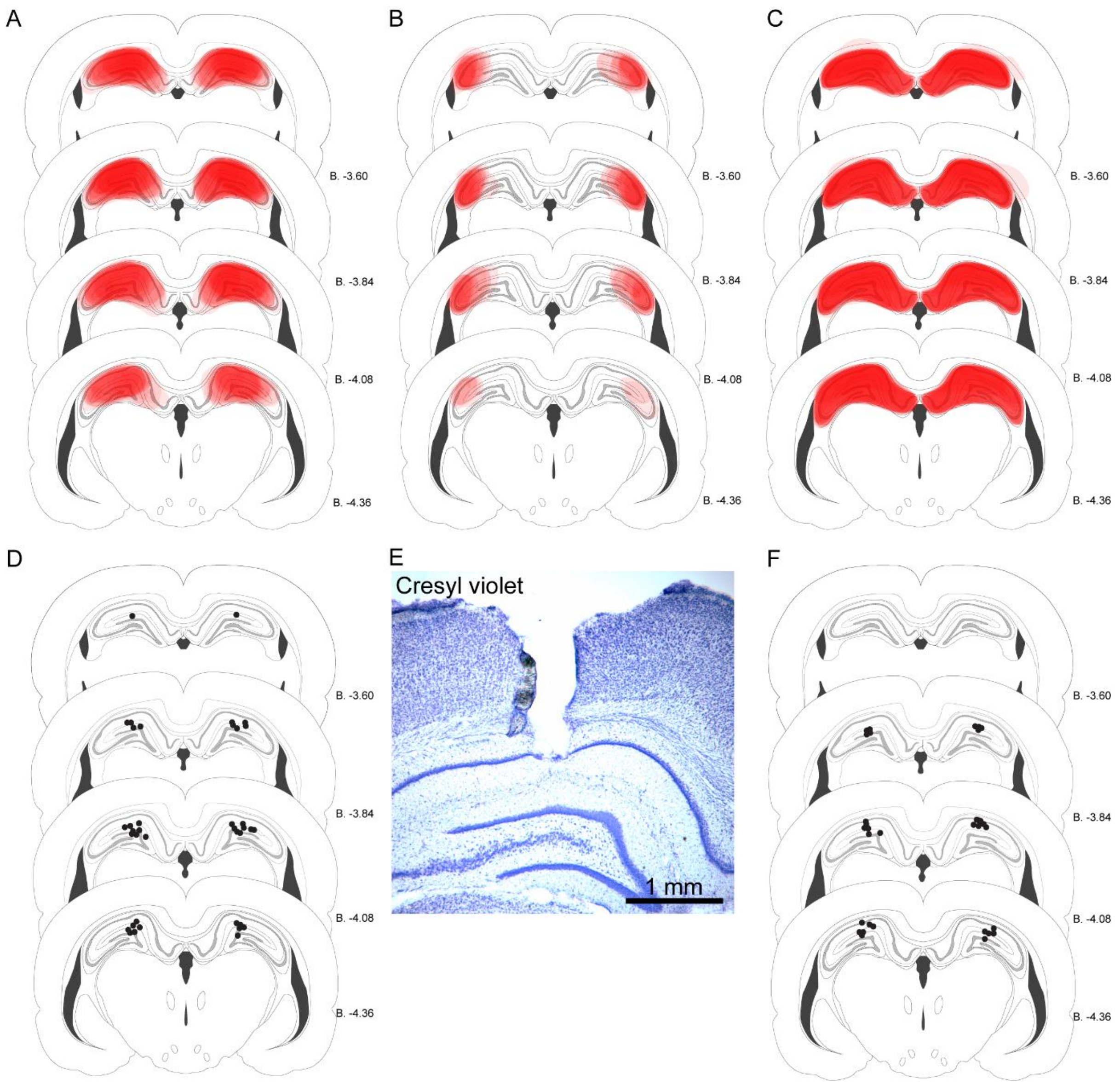


**Figure S3. Cannula and DREADDs placements.** A) Diagrammatic representation of CA1/Dorsal hippocampal viral DREADDs placements (indicated in red), showing each overlapping placement for Experiment 3. B) Diagrammatic representation of CA2 viral DREADDs placements (indicated in red), showing each overlapping placement for Experiment 3. C) Diagrammatic representation of dorsal hippocampal viral DREADDs placements (indicated in red), showing each overlapping placement for Experiment 4. D) Diagrammatic representation of cannula placements from Experiment S1a (indicated by black dots). E) Photomicrograph of representative cannula placement. F) Diagrammatic representation of cannula placements from Experiment 3b (indicated by black dots). All placements are shown on coronal sections adapted from the rat brain atlas of Paxinos & Watson[5].
